# Semicircular canals in *Anolis* lizards: ecomorphological convergence and ecomorph affinities of fossil species

**DOI:** 10.1101/121525

**Authors:** Blake V. Dickson, Emma Sherratt, Jonathan B. Losos, Stephanie E. Pierce

## Abstract

*Anolis* lizards are a model system for the study of adaptive radiation and convergent evolution. Greater Antillean anoles have repeatedly evolved six similar forms or ecomorphs: crown-giant, grass-bush, twig, trunk, trunk-crown, and trunk-ground. Members of each ecomorph category possess a specific set of morphological, ecological and behavioural characteristics which have been acquired convergently. Here we test whether the semicircular canal system – the organ of balance – is also convergent among ecomorphs, reflecting the shared sensory requirements of their ecological niches. As semicircular canal shape has been shown to reflect different locomotor strategies, we hypothesised that each *Anolis* ecomorph would have a unique canal morphology. Using 3D semilandmarks and geometric morphometrics, semicircular canal shape was characterised in 41 *Anolis* species from the Greater Antilles and the relationship between canal shape and ecomorph grouping, phylogenetic history, size, and perch characteristics was assessed. Further, canal morphology of modern species was used to predict the ecomorph affinity of five fossil anoles from the Miocene of the Dominican Republic. Our study recovered ecomorph as the single-most important covariate of canal morphology in modern taxa; although phylogenetic history and size also showed a small, yet significant correlation with shape. Surprisingly, perch characteristics were not found to be significant covariates of canal shape, even though they are important habitat variables. Using posterior probabilities, we found that the fossil anoles have different semicircular canals shapes to modern ecomorph groupings implying extinct anoles may have been interacting with their Miocene environment in different ways to modern *Anolis* species.

## 1. Introduction

The semicircular canals are a functional component of the vestibular system of the inner ear that enable vertebrate animals to coordinate fast and complex movements in three-dimensions (3D). As such, it stands to reason that more agile and mobile animals, such as fast moving arboreal species, would benefit from an enhanced sense of balance, granted through refinement of canal morphology. Recent theoretical [1,2], physiological [3,4], and experimental [5–9] studies have shown that in mammals there is a relationship between locomotor performance – an important component of ecological niche – and canal morphology. Although these studies use different metrics for canal morphology, such as canal size [7,10], torsion [5] and orthogonality [2,6], all agree there is a strong signal between semicircular canal morphology and locomotor ability. Thus, our current understanding of the vestibular system in living mammals has allowed us to investigate and interpret the behaviour and ecology of extinct species [11–21].

The vestibular system has, however, been little explored outside the Mammalia. While the semicircular canals are physiologically and anatomically homologous in all vertebrates, we cannot assume that the relationship between form and function in mammals will hold true for other taxa. Extrapolation from mammals is particularly problematic for studies wishing to reconstruct the ecology of non-mammalian fossil species [20,22–25]. Recent studies looking at the morphology of the semicircular canals with respect to ecology in amphibians [26], squamates [27,28], and birds [29,30] have begun to expand our knowledge of the vestibular system beyond mammals, though much work is still needed to fully understand the system from a greater evolutionary and ecological spectrum.

*Anolis* lizards (Dactyoidae) represent a unique opportunity for furthering such research. Greater Antillean anoles encompass a high level of phylogenetic diversity [31,32], but their morphology is strongly constrained by the requirements of the niches they inhabit. Within *Anolis*, six ecomorph types have evolved independently on each of the four Greater Antilles islands (with several exceptions), with each ecomorph encompassing a specific suite of anatomical (e.g. short vs. long limbs), ecological (e.g. tree trunk vs. branches), and behavioural (e.g. locomotion, territoriality) characteristics (reviewed in [33]). The recurrent and consistent adaptive radiations found in *Anolis* make it an excellent starting point for further understanding the morpho-functional relationship of the semicircular canals in squamates and beyond Mammalia in general.

Here, we use 3D geometric morphometrics to quantify and investigate whether *Anolis* semicircular canal morphology is convergent among ecomorph groups. Specifically, we aim to test the hypothesis that anole species adapted to similar ecomorph niches have converged on similar semicircular canal morphologies, reflecting the sensory requirements of their shared ecological and behavioural habits. We also test the influence of phylogenetic relatedness and size on patterns of canal morphology, two factors that have been shown to have an effect on vestibular system form [7,9,27,34–38], as well as perch height and diameter – the two most frequently reported habitat variables [33]. Further, we reconstruct the vestibular system in five 15-20 Ma fossil anole specimens preserved in Miocene amber [39] and use our extant dataset to predict the ecomorph affinities and paleoecology of these extinct *Anolis* lizards.

## 2. Material and methods

### Specimens and sample size

The sample consists of 131 individuals representing 41 species of anoles originating from the four islands of the Greater Antilles: Hispaniola, Cuba, Jamaica and Puerto Rico. Species from Hispaniola are represented by multiple specimens including juvenile individuals (see further below). All six ecomorphs are represented by multiple species: crown-giant (5), grass-bush (6), trunk (3), trunk-crown (9) trunk-ground (4) and twig (5), with the addition of eight ‘unique’ species that are endemic to each island but do not conform morphologically to any of the specified ecomorphs [33]. Five fossil anoles of Miocene age (Figure 1) were also included from the amber deposits of the Dominican Republic [39,40], details of which can be found in Sherratt *et al*. [39]. All modern specimens were sourced from the Herpetology collection at the Museum of Comparative Zoology (MCZ), Harvard University. All species and specimen numbers can be found in Table S1.

**Figure 1.**
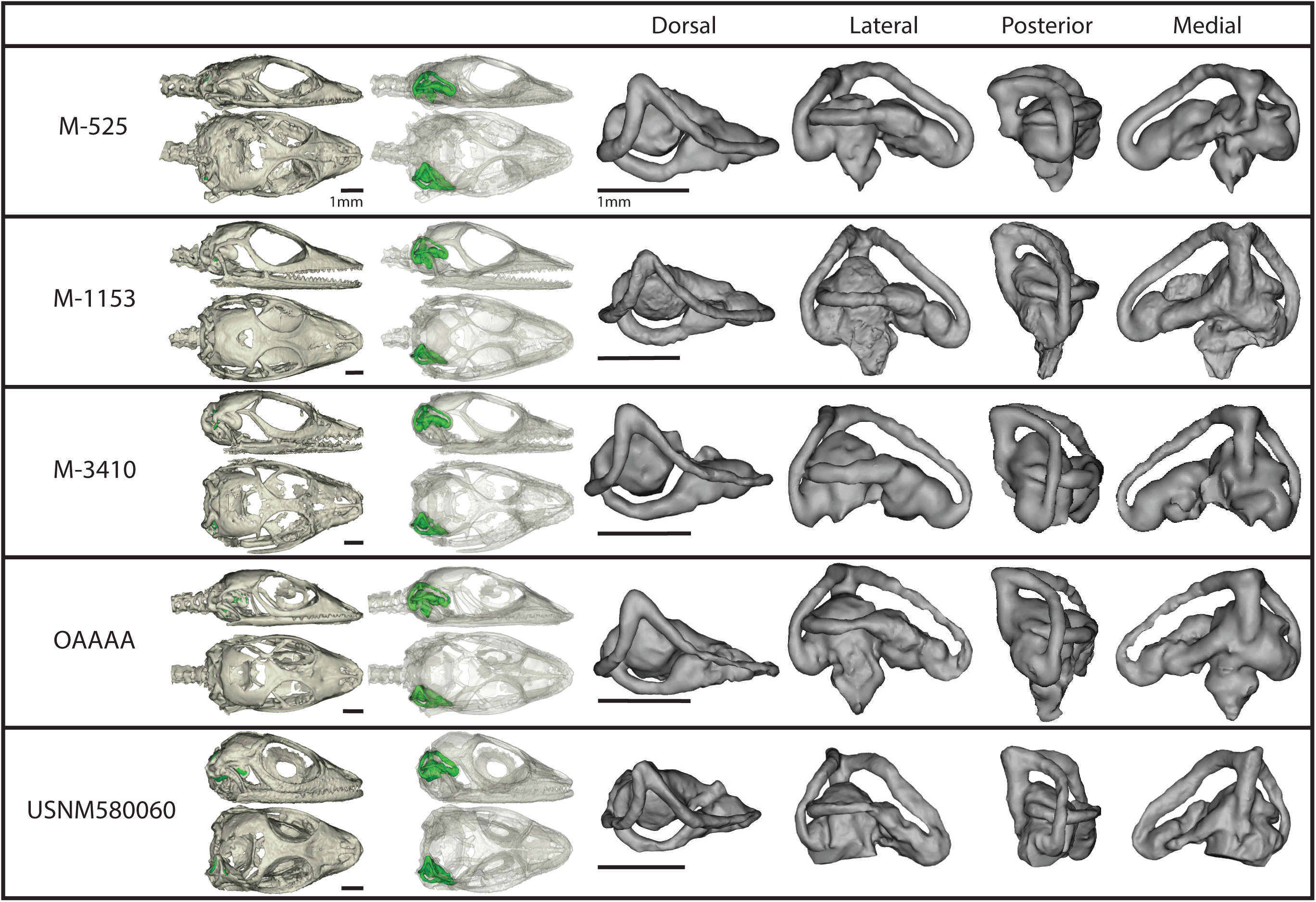
3D rendering of the amber fossil specimens showing the vestibular system within the skull, and the vestibular system in anterior, posterior, lateral, and dorsal views. Scale bar = 1mm.

### Data Acquisition and Landmarks

Various methods for measuring the complex structure of the three semicircular canals have been used to date. Traditional morphometric approaches, in the form of linear and angular measures of size and orthogonality, have been used extensively in the past [2,6,7,10] with the benefit of being easily comparable across studies. However, these measurements struggle to completely capture the full shape variation in the canals owing to their complex curvature. Geometric morphometrics (GM; [41,42]) has been used increasingly to overcome this shortcoming, with landmark [9,15,27,34] and semilandmark [18,43,44] approaches, particularly the latter, capable of capturing far more morphological variation than standard morphometrics. GM methods do, however, vary between studies and are thus less easily compared across studies and broader taxonomic groups. For morphometric approaches, digital thresholding and segmenting of micro computed tomography (μCT) data can introduce significant variation in canal lumen thickness [45], making measuring and digitizing the canal surface error prone. Instead, utilizing a centreline through the lumen overcomes this potential error as it is not effected by threshold values [43,45]. This is the approach that we took.

Specimens were μCT scanned using a variety of imaging systems and settings (Table S1) and the semicircular canals manually segmented and 3D rendered using Materialise Mimics® software. 3D landmarks were derived from the centreline of the semicircular canals, calculated using the ‘medcad’ module in Mimics®, and then manually adjusted to optimise the position of the centreline through the lumen. This centreline was then split into four segments – the three canals (anterior, posterior and lateral) and the crus commune (Figure 2a) – and exported as 3D coordinates describing a curve. The centreline curves were then resampled using the Resample executable [46] so that the three canals were each described by 28 equally-spaced semilandmarks, and the crus commune by three semilandmarks. These semilandmarks were anchored by five landmarks positioned at the junction of the anterior and lateral canals, the posterior and lateral canals with the vestibule, and the junction of the crus commune, resulting in 95 landmarks in total (Figure 2b).

**Figure 2.**
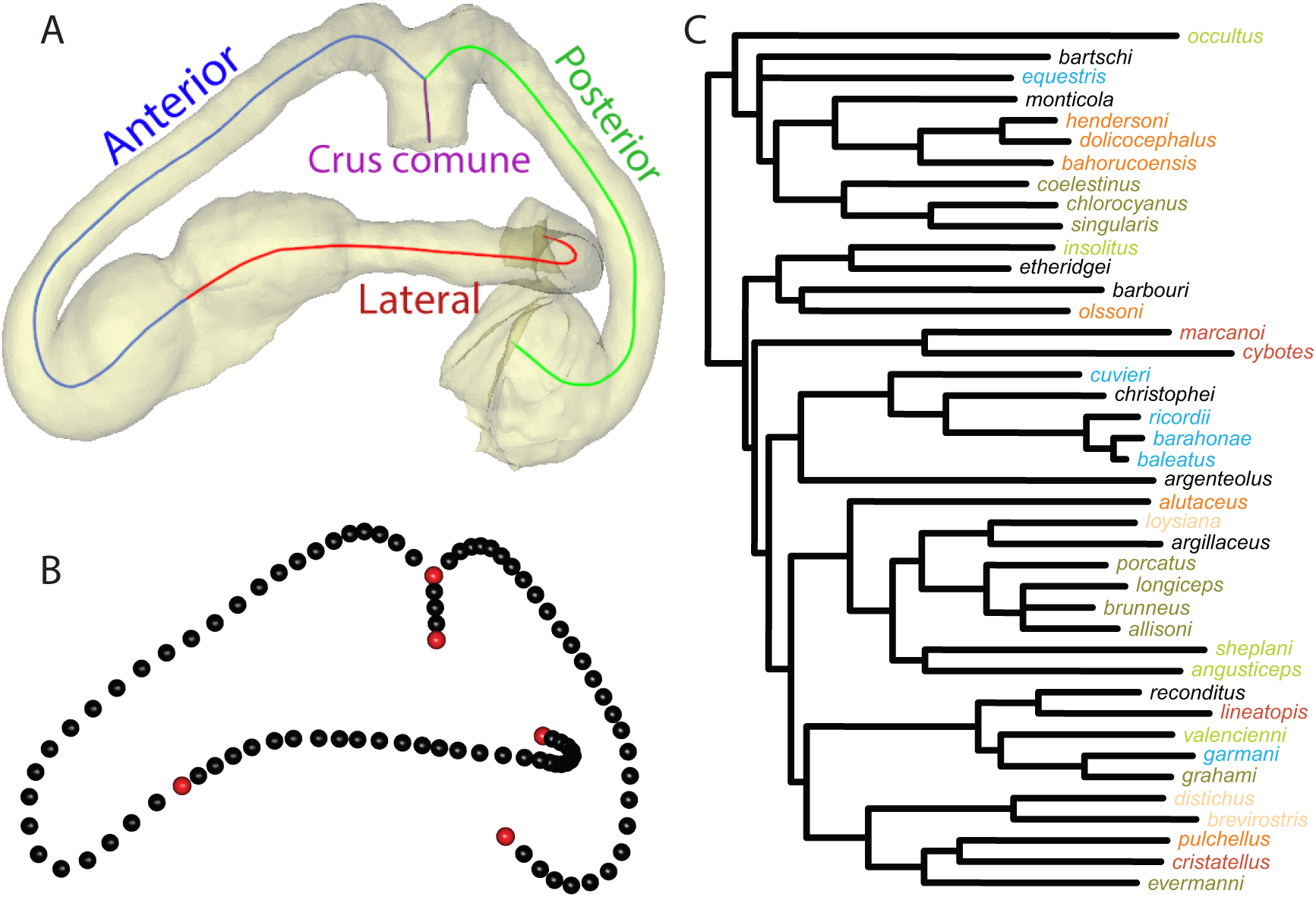
Segmented semicircular canal demonstrating placement of (A) centrelines and (B) semilandmarks; and (C) a time calibrated phylogeny of study taxa coloured by ecomorph. Red landmarks are fixed, while black landmarks are sliding. Species colour coding: Blue=crown-giant; Orange-grass-bush; Pink=trunk; Army Green=trunk-crown; Red=trunk-ground; Bright Green=twig; Black=unique.

### Data Analysis

Landmark coordinate data were aligned by Procrustes superimposition, allowing semilandmarks to slide along their tangent directions in order to minimise bending energy [47,48] using the R statistical environment v.3.2.3 [49] and the package *geomorph* v.2.1.4 [50]. The resulting Procrustes residuals were used as shape variables in the subsequent analyses. Alignment was done on both the full shape variable dataset (n=131) and a phylogenetic subset (n=41). Since the full dataset represented ontogenetic series of each species, the phylogenetic subset was represented by the largest adult individual from each species paired with the phylogeny of Gamble *et al*. [51] (Figure 2c). Partitioning the data was necessary to investigate the role phylogeny might play in determining canal shape as no current methods are available that would account for ontogenetic variation in phylogenetic comparative analyses.

Principal component analysis (PCA) was performed on the phylogenetic subset dataset to visualise semicircular canal shape variation among all species. Eigendecomposition of this PCA was performed using only modern taxa (including unique species); fossil specimens were later projected into this morphospace using the PCA eigenvectors. The phylogeny [51] was projected into the phylogenetic subset PC space to visualise the estimated evolutionary trajectory of canal shape change, using the *geomorph* function ‘plotGMPhyloMorphoSpace’. Phylogenetic signal of canal morphology was calculated using the K statistic [52,53] with *geomorph*’s ‘physignal’ function and tested for significance using 10,000 permutations. To visualise a morphospace independent of the effects of phylogeny and allometry, we plotted the residuals of a phylogenetic regression with log-transformed semicircular canal centroid size, performed on PC scores using the ‘phyl.resid’ function of *phytools* v.0.5-10 [54,55]. Centroid size is a measure of size calculated as the square root of the sum of squared distances of a set of landmarks from their centroid [42].

To determine whether ecomorphs occupy different regions of morphospace (and thus have significantly different canal morphologies), and to test the effect of size on canal shape, analysis of covariance (ANCOVA) and phylogenetic generalised least squares (PGLS) were performed using the ‘procD.lm’ and ‘procD.pgls’ functions respectively [56] of *geomorph*. These functions perform statistical assessment of the terms in the model using Procrustes distances among specimens, rather than explained covariance matrices among variables, and are thus suitable for multivariate datasets [56,57]. Three log-transformed size metrics were used: semicircular canal centroid size, skull length (measured between the premaxilla and the dorsal margin of the foramen magnum) and skull width (measured between the paraoccipital processes). The relationship between the size metrics was also explored using linear regressions. In addition, we investigated the relationship between canal shape and perch height and diameter ([50] and unpubl.) by ANCOVA and PGLS.

Finally, a canonical variate analysis (CVA) was run using the “CVA” function of the package *Morpho* v.2.3.0 [58] to explore the morphological shape variables that maximise between ecomorph group variance relative to within group variance, and to predict the potential ecology of the fossil anoles. Prior to running the CVA, a PCA was performed on the full extant dataset (excluding unique species) and the first 40 PC axes representing 99% of the variation were extracted; this reduction in dimensionality was done to ensure that the number of shape variables (n = 40) was less than the number of individual specimens (n=99) [59] and to remove minor components of shape variance that might be attributable to error. In addition, the full specimen dataset was used to take into consideration both specific and ontogenetic variation and to increase the power of the test by incorporating a larger sample size. Within the CVA morphospace, 95% confidence intervals were generated around each of the modern ecomorph groups. The unique and fossil specimens were then projected into this morphospace using the canonical variates. To establish whether fossil specimens and unique taxa fall within the morphological ranges of modern ecomorphs, Mahalanobis distances were calculated between fossils, unique species and each ecomorph group, and compared to within-group variations [60]. Probabilities were then calculated non-parametrically using resampling of within-group variation with 10,000 replicates. Log-likelihood estimations were also calculated to allow comparison with previous work [39].

## 3. Results

### Patterns of shape variation

PCA (Figure 3a, b) shows that PC1 (40.2% of variation) largely represents changes in anterior and lateral canal morphology. Moving from PC1 positive to PC1 negative, there is a trend for the canals to become more rounded and anterodorsally shortened. PC2 (11.0% of variation) represents moderate changes in all three canals, with a PC2 positive to PC2 negative shift showing rounding of the anterior-most section of the anterior canal, more torsion (out-of-plane curvature) of the lateral canal, and less torsion of the posterior canal. PC3 (10.5% of variation) represents changes in the anterior and posterior canals, with a transition from PC3 positive to PC3 negative showing increased curvature and deepening of the posterior canal and reduction of the lateral aspect of the anterior canal.

**Figure 3.**
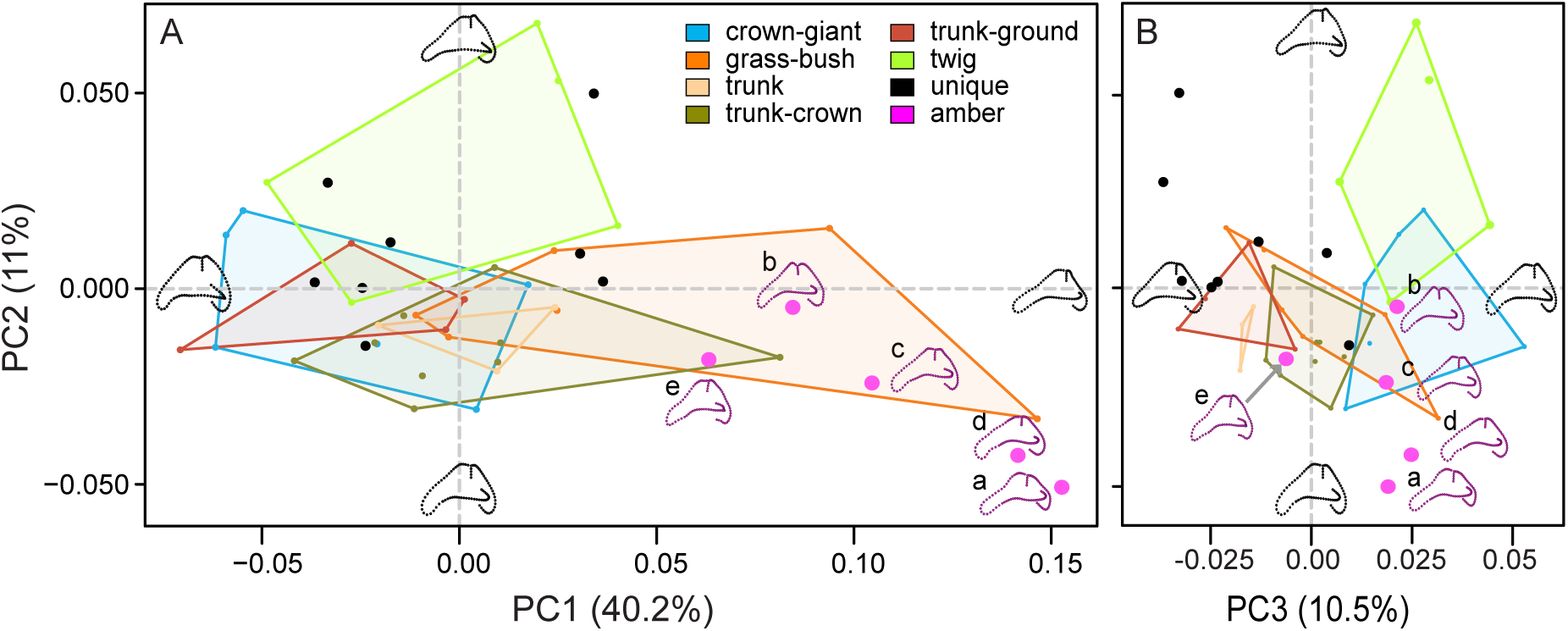
Principal component (PC) analysis of semicircular canal shape showing (A) PC1 vs. PC2 and (B) PC2 vs. PC3. The first three PCs represent 61.7% of variation in canal shape. Points are specimens, coloured by ecomorph and bounded by convex hulls. Unique specimens are shown in black, and fossil specimens in magenta. Thin-plate spline (TPS) warps representing the maxima and minima of each PC are shown on each axis, and TPS warps of the fossils are shown in magenta. Amber fossils: a= M-525, b= M-1153, c= M-3410, d= OAAAA, e= USNM580060.

Visually, there is significant overlap between ecomorph groupings along PC1 and PC2, with grass-bush anoles occupying most of PC1. PC3 separates the three ‘trunk’ ecomorphs from the twig and crown-giant ecomorphs. When the shape data were corrected for size and phylogeny, morphospace is minimally altered (Figure S1). The unique species are widely distributed across morphospace, overlapping with most ecomorph grouping (PC2 vs. PC1) and falling outside the variation enclosed by the ecomorphs (PC2 vs. PC3). All fossil specimens fall along the positive end of PC1, in the grass-bush area of morphospace, which represents flattening and anterodorsal elongation of the anterior canal. Furthermore, three fossil specimens (M-1153, M-3410, USNM580060) overlap with multiple ecomorph groupings along PC3. Fossil specimens M-525 and OAAAA appear to fall beyond the morphologies of all living taxa sampled.

### Predictors of shape

Phylogeny was found to have only a weak influence on semicircular canal morphology (K=0.58), though permutation found this influence to be greater than expected from random (P=0.0034). Mapping of the phylogeny onto morphospace (Figure 4) shows extensive overlapping of branches through morphospace, indicating convergence towards similar semicircular canal morphologies.

**Figure 4.**
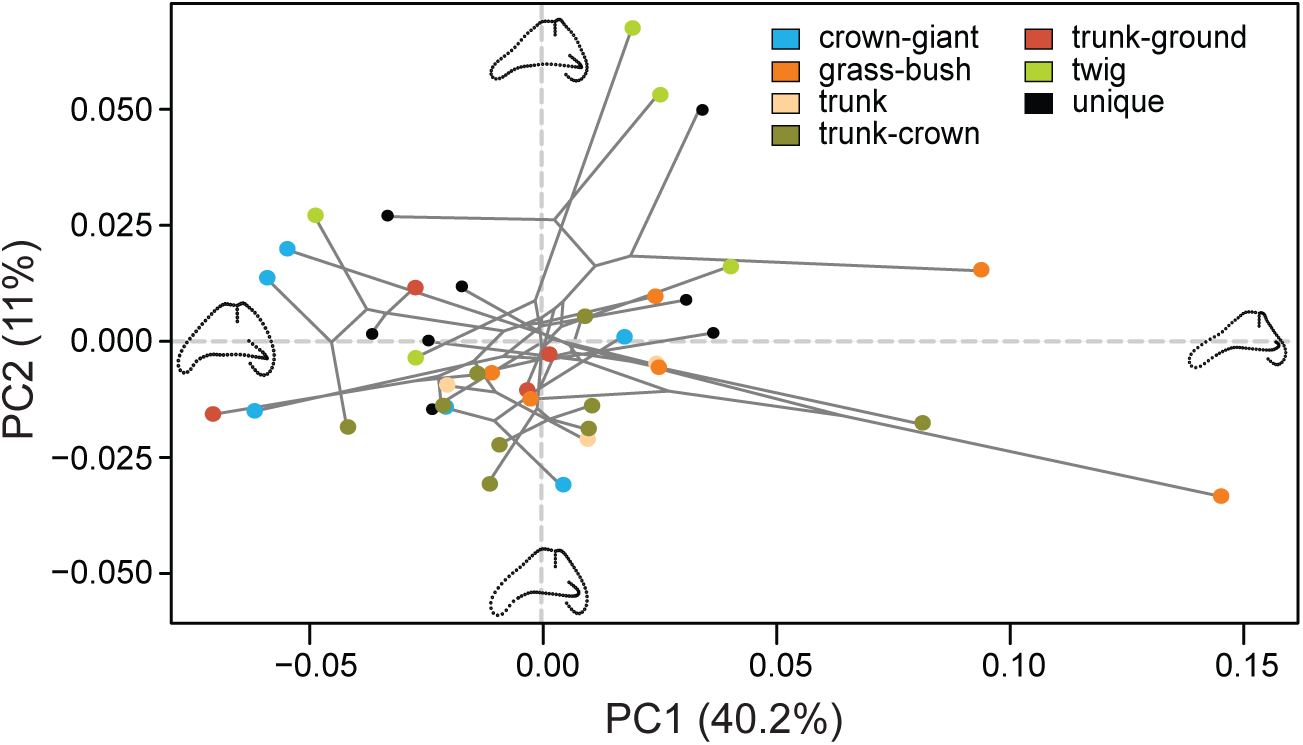
Time calibrated *Anolis* phylogeny projected into the morphospace of PC1 vs. PC2, representing 51.2% of the total variation. Points are coloured by ecomorph, with unique species in black. TPS warps representing the maxima and minima of each PC are shown in black on each axis.

ANCOVA (Table 1) found that ecomorph is a strong and significant predictor (R^2^=0.35, P<0.001) of canal shape. A weak but significant relationship also exists between centroid size and canal shape (R^2^=0.09, P=0.007). Further, PGLS (Table 1) found ecomorph to be a significant predictor of canal shape, though the effect was less strong (R^2^=0.25, P<0.001). This reduction in the correlation coefficient indicates an interaction between phylogeny and ecomorph and that ecomorph groupings are not entirely independent of phylogeny. PGLS (Table 1) also revealed a significant relationship between centroid size and canal shape (R^2^=0.13, P<0.001) and a significant interaction between ecomorph and centroid size (R^2^=0.18, P=0.037). Similar results were recovered when using skull length and width as a measure of size (Table S2), both of which are strongly related to centroid size (linear regression, R^2^= 0.93, P<0.001; R^2^=0.89, P<0.001 respectively). There was no significant relationship between either perch height or diameter and canal shape, with and without phylogenetic correction (Table 1).

**Table 1.**
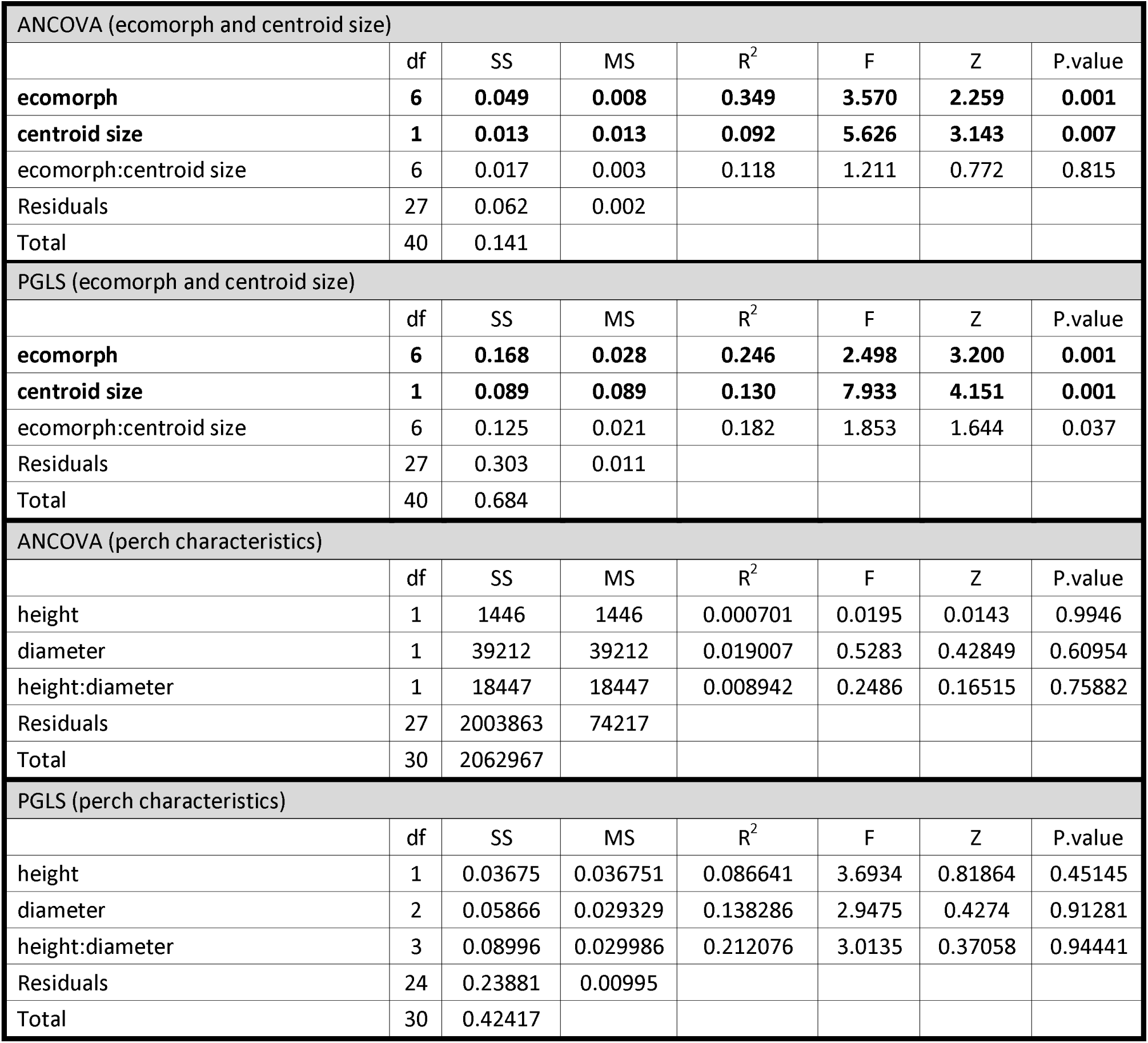
Analysis of covariance (ANCOVA) and phylogenetic generalised least squares (PGLS) of semicircular canal shape against ecomorph, semicircular canal centroid size, and perch characteristics with statistical significance assessed through 10,000 permutations.

| ANCOVA (ecomorph and centroid size) |  |  |  |  |  |  |  |
| --- | --- | --- | --- | --- | --- | --- | --- |
|  | df | SS | MS | R<sup>2</sup> | F | Z | P.value |
| **ecomorph** | **6** | **0.049** | **0.008** | **0.349** | **3.570** | **2.259** | **0.001** |
| **centroid size** | **1** | **0.013** | **0.013** | **0.092** | **5.626** | **3.143** | **0.007** |
| ecomorph:centroid size | 6 | 0.017 | 0.003 | 0.118 | 1.211 | 0.772 | 0.815 |
| Residuals | 27 | 0.062 | 0.002 |  |  |  |  |
| Total | 40 | 0.141 |  |  |  |  |  |
| PGLS (ecomorph and centroid size) |  |  |  |  |  |  |  |
|  | df | SS | MS | R<sup>2</sup> | F | Z | P.value |
| **ecomorph** | **6** | **0.168** | **0.028** | **0.246** | **2.498** | **3.200** | **0.001** |
| **centroid size** | **1** | **0.089** | **0.089** | **0.130** | **7.933** | **4.151** | **0.001** |
| ecomorph:centroid size | 6 | 0.125 | 0.021 | 0.182 | 1.853 | 1.644 | 0.037 |
| Residuals | 27 | 0.303 | 0.011 |  |  |  |  |
| Total | 40 | 0.684 |  |  |  |  |  |
| ANCOVA (perch characteristics) |  |  |  |  |  |  |  |
|  | df | SS | MS | R<sup>2</sup> | F | Z | P.value |
| height | 1 | 1446 | 1446 | 0.000701 | 0.0195 | 0.0143 | 0.9946 |
| diameter | 1 | 39212 | 39212 | 0.019007 | 0.5283 | 0.42849 | 0.60954 |
| height:diameter | 1 | 18447 | 18447 | 0.008942 | 0.2486 | 0.16515 | 0.75882 |
| Residuals | 27 | 2003863 | 74217 |  |  |  |  |
| Total | 30 | 2062967 |  |  |  |  |  |
| PGLS (perch characteristics) |  |  |  |  |  |  |  |
|  | df | SS | MS | R<sup>2</sup> | F | Z | P.value |
| height | 1 | 0.03675 | 0.036751 | 0.086641 | 3.6934 | 0.81864 | 0.45145 |
| diameter | 2 | 0.05866 | 0.029329 | 0.138286 | 2.9475 | 0.4274 | 0.91281 |
| height:diameter | 3 | 0.08996 | 0.029986 | 0.212076 | 3.0135 | 0.37058 | 0.94441 |
| Residuals | 24 | 0.23881 | 0.00995 |  |  |  |  |
| Total | 30 | 0.42417 |  |  |  |  |  |

### Ecomorph Differences

The CVA biplots (Figure 5) and posterior probabilities (Table 2) show ecomorphs form significantly different groups within the morphospace (P<0.0001). On the extreme negative end of canonical function 1 (CF1, Figure 5a) are the twig and crown-giant ecomorphs. This region of morphospace is characterized by greater out-of-plane curvature of the anterior canal. The two groups, however, occupy opposite extremes of CF2 (Figure 5a, b). Crown-giants occupy the extreme positive end of CF2 and, when compared with twig species, display greater dorsoventral curvature and reduced torsion of the posterior canal, and lengthening of the crus-commune. Grass-bush and the three ‘trunk’ ecomorphs occupy the central and positive region of CF1, characterised by reduced anterior canal curvature, but they do separate along CF2 and CF3 (Figure 5b). Trunk-crown and grass-bush species occupy a more positive region along CF2, displaying greater dorsoventral curvature of the anterior canal, reduced torsion of the posterior canal, and longer crus-commune. Trunk-crown and grass-bush species separate along CF3 with the trunk-crown ecomorph occupying the positive end of CF3 indicating greater curvature of the lateral canal and reduced curvature of the posterior canal. The remaining ‘trunk’ ecomorphs separate along CF2, with the trunk-ground species occupying a more negative region of CF2, representing reduced torsion of the canals in general.

**Figure 5.**
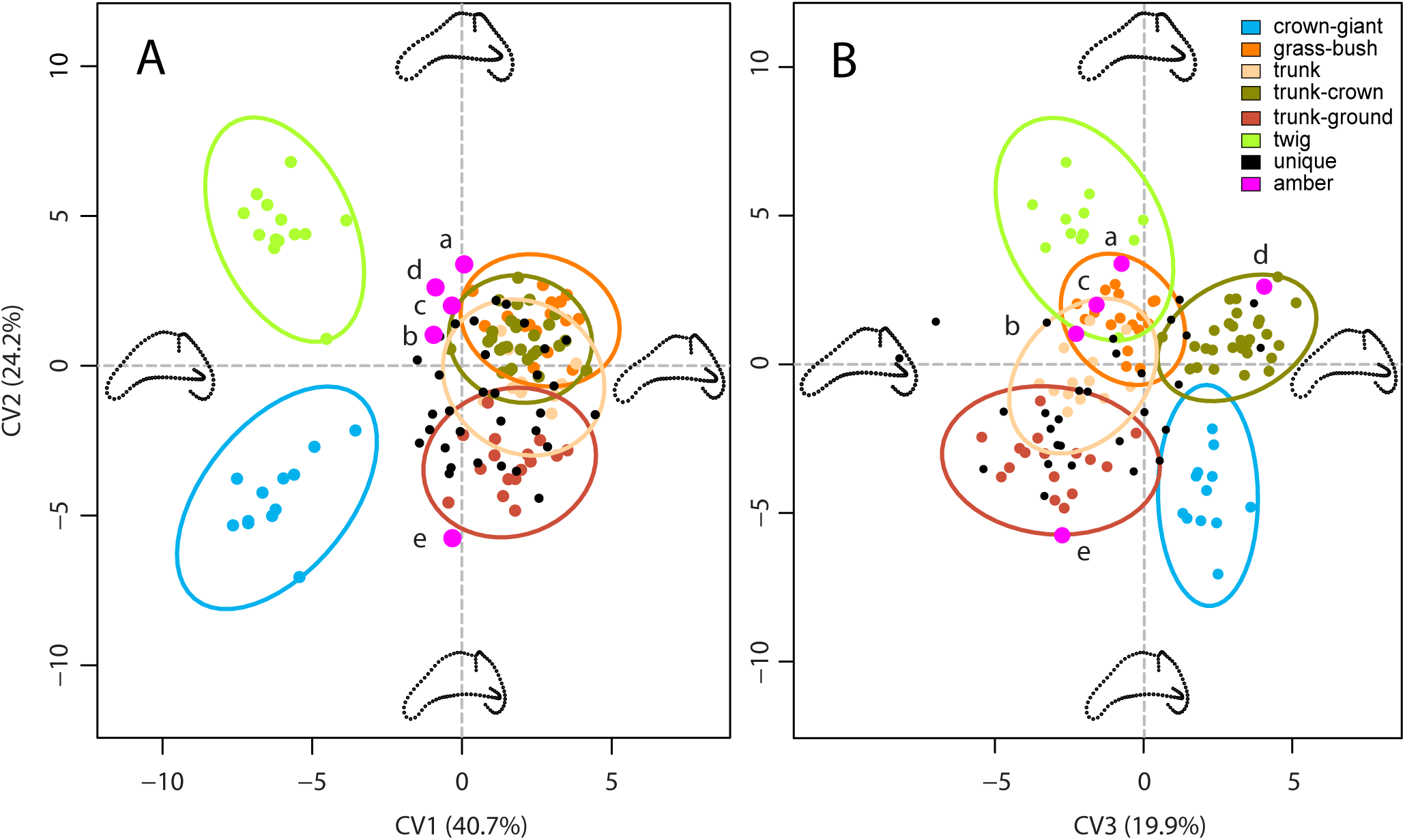
Canonical variates (CV) analysis showing (A) CF1 vs. CF2, and (B) CF2 vs. CF3, representing 84.8% of the total variation. 95% confidence ellipses of each ecomorph are plotted, and points are coloured by ecomorph, with unique species plotted in black and fossils specimens in magenta. Thin-plate spine (TPS) warps representing the maxima and minima of each CF are shown in black along each axis. Amber fossils: a= M-525, b= M-1153, c= M-3410, d=amber OAAAA, e= USNM580060.

**Table 2.** Posterior probabilities of ecomorph groups being significantly different from one another based on Mahalanobis distances using 10,000 permutations. CG=crown-giant, GB=grass-bush, TC=trunk-crown, TG=trunk-ground, Tr=trunk,Tw=twig.

| Probabilities from Mahalanobis Distances |  |  |  |  |  |  |
| --- | --- | --- | --- | --- | --- | --- |
|  | CG | GB | TC | TG | Tr | Tw |
| CG |  | <0.0001 | <0.0001 | <0.0001 | <0.0001 | <0.0001 |
| GB | <0.0001 |  | <0.0001 | <0.0001 | <0.0001 | <0.0001 |
| TC | <0.0001 | <0.0001 |  | <0.0001 | <0.0001 | <0.0001 |
| TG | <0.0001 | <0.0001 | <0.0001 |  | <0.0001 | <0.0001 |
| Tr | <0.0001 | <0.0001 | <0.0001 | <0.0001 |  | <0.0001 |
| Tw | <0.0001 | <0.0001 | <0.0001 | <0.0001 | <0.0001 |  |

The unique species (those not belonging to any of the ecomorph categories) are spread throughout morphospace (Figure 5), with only some visually falling within the 95% CI of modern ecomorphs in the first few axes (CF1-3). Posterior probabilities of Mahalanobis distances over all CFs find that many unique species fall significantly outside the 95% CIs defined by the ecomorphs, with exceptions (Table S3): 1) one specimen of *A. argenteolus* falls within the 95% CI of trunk-crown (P=0.139); 2) the sole *A. bartschi* specimen falls within the 95% CI of trunk-crown (P=0.895); 3) out of five *A. christophei* specimens, one falls within the 95% CI of trunk-ground (P=0.223), one in trunk-crown (P=0.151), and one in grass-bush (P=0.073); 4) out of five *A. etheridgei* specimens, two fall within the 95% CI of trunk-ground (P=0.191, P=0.052); 5) out of five *A. montícola* specimens, three fall within the 95% CI of trunk-ground (P=0.209, P=0.075, P=0.348) and one in trunk (P=0.103); 6) the only *A. reconditus* falls within trunk-ground (P=0.124); and 7) out of four *A. rimarum* specimens, one falls within the 95% CI of grass-bush (P=0.478) and one in trunk-crown (P=0.059). Our log-likelihood calculations do, however, assign most unique species to the defined ecomorphs, although these assignments are also inconsistent within and between species (Table S3).

All the fossil specimens fall either outside or just on the margin of the 95% CI of the modern ecomorphs, much like the unique species (Figure 5). Our posterior probabilities support this: all fossil specimens are highly unlikely to belong to any modern ecomorph group (Table 3). This contrasts with the log-likelihood tests which assign each fossil to the ‘closest’ ecomorph group regardless of actual morphological distance (Table 3). Moreover, log-likelihood tests found only one instance of correspondence with the ecomorphs inferred by Sherratt *et al*. [39]: OAAAA, which is assigned to trunk-crown. Visually, all fossils except USNM580060 fall into an intermediate region of the CVA morphospace, suggesting a generalised lifestyle. USNM580060 falls into the negative region of CF2, closest in morphology to trunk-ground ecomorphs, though it falls outside the trunk-ground 95% CI (Table 3).

**Table 3.** Probabilities and log-likelihoods of fossil anoles belonging to modern ecomorph groups, compared with the results of Sherratt *et al*. [39]. Probabilities below 0.05 indicate the fossil is significantly different from an ecomorph group. Likelihoods are calculated as the most likely modern group without an alternative hypothesis (i.e. fossils are unique). CG=crown-giant, GB=grass-bush, TC=trunk-crown, TG=trunk-ground, Tr=trunk, Tw=twig.

|  | Log-Likelihood | Probability |  |  |  |  |  | Sherratt *et al.* [39]<br>Log-Likelihood |
| --- | --- | --- | --- | --- | --- | --- | --- | --- |
|  |  | CG | GB | TC | TG | Tr | Tw |  |
| **M-1153** | **GB (0.99)** | <0.0001 | 0.0313 | <0.0001 | <0.0001 | <0.0001 | <0.0001 | **TC (1.00)** |
| **M-3410** | **GB (0.99)** | <0.0001 | 0.0108 | <0.0001 | <0.0001 | <0.0001 | <0.0001 | **TC (0.69)** |
| **M-525** | **TW (0.90)** | <0.0001 | <0.0001 | 0.0002 | <0.0001 | <0.0001 | 0.0018 | **TC (0.99)** |
| **OAAAA** | **TC (0.99)** | <0.0001 | <0.0001 | 0.0086 | <0.0001 | <0.0001 | <0.0001 | **TC (1.00)** |
| **USNM580060** | **CG (0.21)** | <0.0001 | <0.0001 | <0.0001 | <0.0001 | <0.0001 | <0.0001 | **TG (0.99)** |

## 4. Discussion

### Convergence of semicircular canal shape

Our results support the hypothesis that phylogenetically disparate *Anolis* species have convergent semicircular canal morphologies, allowing them to navigate similar ecological niches. We found ecomorph grouping is by far the strongest determinant of canal shape, even when phylogeny is accounted for (25-35%, Table 1). Given the role semicircular canals play in coordinating fast and complex movements in three-dimensions, and the differences in ecology and locomotor behaviour among the *Anolis* ecomorphs, this result aligns well with the body of work supporting a relationship between semicircular canal morphology and locomotion [5,7,9,21,26,27].

Though differences among the ecomorphs are subtle, both crown-giant and twig ecomorphs display more torsion of the anterior canal than those of the other ecomorphs, separating significantly along CF1 (Figure 5a). Increasing out-of-plane torsion of a canal may increase sensitivity to rotations out of the canal’s major plane of motion detection [61]. Further, CF3 resolves trunk-crown and crown-giant ecomorphs based on increased curvature and length of the lateral canal (Figure 5). Increasing canal length suggests greater sensitivity [3]. These potential increases in sensitivity of both anterior and lateral canals may represent adaptations to the specialised arboreal niches of these three ecomorphs – crown-giant, twig and trunk crown – occupying the complex upper reaches of the canopy, requiring greater sensitivity to movements. For the remaining ecomorphs, out of plane sensitivity may not be as essential to locomotor performance. For trunk and trunk-ground ecomorphs the trunk provides a broad surface on which locomotion is much easier [62–64] requiring less refined balance. The perch diameter for grass-bush ecomorphs is indeed relatively much more narrow [33,65], however, the consequences of falling from grass or a bush are far less severe than falling from the tree canopy as in the higher dwelling ecomorphs.

Generally, however, the ecomorphological signal we found does not fully explain variation in semicircular canal shape (Table 1). Analysis of canal shape and perch height and diameter returned non-significant results (Table 1), despite both being correlated with ecomorph [33]. This finding was unexpected given the importance of balance during locomotion on narrow perches, and the assumed consequences of falling from high perches. The perch data included here are from a different population than our morphological data; perhaps this limitation introduced sufficient error into our analysis to confound the relationship. Further work is needed. Other behavioural characteristics may also be associated with canal shape variation, such as locomotor performance over varied substrates and/or head rotational velocities [18], and we encourage collection of such data. The remaining variation may also include morphological ‘noise’ – with the semicircular canals conforming to the other anatomical requirements of the skull, such as the brain, the feeding apparatus and the other senses of sight and hearing.

### Role of phylogeny and size

We found a small yet significant relationship between phylogenetic relatedness and semicircular canal morphology (Figure 4; Table 1). This weak phylogenetic signal may be the result of repeated convergent evolution for which anoles are famous [33], though similarly weak yet significant phylogenetic signals are consistent across studies dealing with other taxonomic groups [27,34–36]. Based on these significant phylogenetic signals, some authors have suggested using the inner ear as a source of phylogenetic characters [66,67]. However, the results of our study demonstrate that while semicircular canal morphology is related to phylogenetic history, size and ecology are more important factors (Table 1) and any phylogenetic analysis based on such characters would be unreliable. Billet *et al*. [68] concluded similarly in their phylogenetic analysis of litopternan petrosal and inner ear characters, finding a potentially confounding allometric signal.

Skull size (and canal size) is correlated with semicircular canal morphology in *Anolis* and our results also show that it co-varies with ecomorph (Table 1). This interaction suggests differences among the allometric shape trajectories of the six ecomorph groups. Previous studies have found that canal size appears to scale with negative allometry, such that smaller animals have relatively larger canals [7,9,36,37]. Some have postulated that smaller animals experience relatively greater angular accelerations of the head than do large animals [7,9,69] and that canal sensitivity is tied to canal radius, suggesting that larger canals are more sensitive to rotation [3]. Our finding that all three size metrics were highly correlated with canal morphology (Table 1, Table S2), but negative (or positive) allometry cannot be determined when the response variable is multivariate. However, regression of log centroid size on skull length in our data set found evidence of strong negative allometry (slope = 0.63; 95% CI = 0.54-0.71) which is in keeping with prior studies. Further analyses exploring the allometric variation in our data will be the subject of future publications.

### Affinities of fossil anoles

Although further research is needed to determine other factors that may co-vary with semicircular canal morphology, the significant relationship between ecomorph and canal shape in extant *Anolis* species enabled us to explore the paleoecology of fossil taxa. Using posterior probabilities, we find that the semicircular canal shapes of all five fossils are significantly different from modern ecomorph groupings (Table 3), and that all five also differ from each other (Figure 1 and 3). It is not unreasonable for the fossil taxa to differ from modern morphological patterns as the ecological context of Miocene anoles was likely different from the modern Antillean ecosystems. Though little is known about the habitat structure of the Antilles during the Miocene, *Hymenaea protera*, the amber forming tree in which the fossils are contained, is more closely related to the African *H. verrucosa* than the modern Antillean species [40,70]. Differences in floral composition in the Miocene may have influenced how extinct anoles where navigating their island environment, meaning semicircular canal shape may have been under different selective pressures.

Using a log-likelihood approach, the fossil anoles are sorted into modern ecomorph groupings, however, only OAAAA matches the predicted groupings of Sherratt *et al*. [39]. It is possible that the discrepancy between the two studies is a result of taphonomic distortion, an unavoidable factor in fossil specimens. Of the five fossil anoles, two fall into an extreme region of PCA morphospace (Figure 3), which could imply taphonomic distortion. However, close inspection of the fossils finds that only M-525 has any noticeable deformation of the basicranium (lateral compression, Figure 1). Therefore, we do not expect taphonomic distortion to be causing these differences between studies. Alternatively, the discrepancy between our study and Sherratt *et al*. [39] may be due to mosaic evolution: the vestibular system may responded differently to selective pressures than other ecomorphological traits (e.g. limb length, digit length, subdigital lamellae) resulting in varying rates of morphological change [71].

Nonetheless, our study highlights a large discrepancy between assigning fossils using probability and likelihood methods. Posterior probabilities are an unrestricted metric to classification, allowing unknown specimens to belong to a group outside of the pre-defined reference samples [60]. In other words, probabilities allow for the alternative hypothesis of the fossil sample not belonging to the observed group data to be considered. Likelihood estimations on the other hand are weighted to make the observed results the most probable given the sample data. The consequence being that likelihood estimations can artificially inflate confidence in group predictions as we demonstrate here by running log-likelihood and probability calculations concurrently (Table 3); for instance, our log-likelihood calculations erroneously sort modern unique species into ecomorph groups with high confidence (Table S3). Taking these statistical differences into consideration, it is more conservative to use the probability approach, particularly when working with fossil samples that are separated from modern species by millions of years or when there is uncertainty about whether an unknown specimen belongs within the known distribution of the sample population.

## 5. Conclusions

Here we demonstrate that the classic ecomorph definitions of *Anolis* of the Greater Antilles are supported by inner ear morphology, with each ecomorph possessing a distinctive semicircular canal shape. We find that ecomorph is the single-most important covariate of canal morphology, although phylogenetic history and canal size are also significantly correlated with canal shape. Surprisingly, we were unable to find any correlation between canal shape and perch variables; however, this result may reflect a mismatch between datasets, as the perch data and morphological data were collected from different populations. Still, much of the morphological variance seen in our sample remains unexplained and further work is required to tease out other ecological, behavioural, and/or anatomical characteristics that may co-vary with semicircular canal morphology. Using the more conservative metric of posterior probabilities, we were unable to assign fossil anoles to modern ecomorph groups. Our results indicate that the semicircular canals of these extinct anoles are morphologically different from modern *Anolis* ecomorphs, suggesting fossil taxa may have been interacting with their Miocene environment in different ways to modern *Anolis* species.

## Acknowledgements

We would like to thank the staff of the MCZ Herpetology collection, specifically José Rosado for access to specimens and scan data; George Lauder for the use of his computers and software; the Morphmet and R online communities; and all the members of the MCZ Vertebrate Paleontology lab: Katrina Jones, Robert Kambic, Phil Lai, Brianna McHorse and Zachary Morris for their help with R and insights during this project.

## Research Ethics

No ethical assessments were required for the conduct of this research

## Animal Ethics

No licences or approvals were required for the conduct of this research

## Permission to carry out fieldwork

No fieldwork was required for the conduct of the research

## Data Availability

All data and code for analyses are available in the Dryad data repository at the following link: http://dx.doi.org/10.5061/dryad.8s586 [72]

## Author Contributions

Concepts and approach were developed by B.V.D and S.E.P, in consultation with J.B.L. CT scan data was collected by E.S and B.V.D. Data analysis and interpretation was performed by B.V.D, E.S., and S.E.P. The manuscript was prepared by B.V.D. and S.E.P., and edited by E.S. and J.B.L. All authors gave final approval for publication.

## Supplementary Information

**Table S1**. List of specimens with collection ID, μCT scan details, ecomorph, island of origin and skull dimensions.

**Figure S1**. Phylogenetic and size corrected principal component analysis (PCA).

**Table S2**. Analysis of covariance (ANCOVA) and phylogenetic generalised least squares (PGLS) of canal shape against ecomorph and skull length and width with 10,000 permutations for significance.

**Table S3**. Probabilities and log-Likelihoods (bold) of unique taxa belonging to an ecomorph.

### Data

Raw data and R script for analyses

